# Detection of the altered metabolites of Raji cells in the presence of Epstein-Barr virus (EBV) using 1HNMR Spectroscopy

**DOI:** 10.1101/314427

**Authors:** Zahra Dindar, Reza Haj Hosseini, Ruhollah Vahhabpoor, Zahra Zamani

**Affiliations:** Department of Biology, Faculty of Basic Science, Payame Noor University, Tehran, Iran; Department of Biology, Faculty of Science, Payame Noor University, Tehran, Iran; Medical Lab Technology Department, School of Allied Medical Sciences, Shahid Beheshti University of Medical Sciences, Tehran, Iran; Biochemistry Department, Pasteur Institute of Iran, Tehran, Iran

**Keywords:** Raji cells, Epstein-Barr virus, 1H NMR Spectroscopy, Metabolites

## Abstract

EBV is one of the most common viruses in humans and is directly implicated in carcinogenesis. The present study aimed to detect the altered metabolites of Raji cells in the presence of Epstein-Barr virus (EBV) using 1H NMR by NOESY technique. The cells and EBV were maintained in RPMI 1640 and 1 ml of cells were transfected into falcon tubes containing cells and were incubated at 37°C for 2 hours. Metabolites extracted using water or chloroform/methanol, lyophilized and sent for 1H NMR analysis by NOESY technique. The NMR spectrum comprising of Fourier Transformed information about metabolites in the control and treated were imported into MATLAB (v.7.8.0.347) software and the metabolic cycles were determined using Metabo-Analyst software. The data have demonstrated that infected cells leads to proliferation and subsequent immortalization of cell lines through cellular replication machinery recruitment and changes the metabolic profile and promotes vital metabolites such as the carbohydrates engage in pentose phosphate and glycolactic, biosynthesis of nucleotide and amino acid pathways. The results also indicate that essential amino acids are required for protecting viral structure and the function of viral genes. Therefore, EBV infection of cells leads to the sustained elevation of cell growth and cell immortalization.

## Introduction

Epstein-Barr virus (EBV) is one of eight known human herpes virus types in the herpes family that was first isolated by Epstein in 1964 from cultured cells of Burkett’s lymphoma frequently found in children of equatorial Africa[1,2]. It is a double-stranded DNA virus of about 170 kb, and encodes about 80 genes. EBV is one of the most common viruses in humans and is directly implicated in carcinogenesis [3] which is associated with the nasopharynx[4], salivary gland [5], breast [6], bladder [7], kidney [8], uterine cervix [9], colon [10] and lung [11] cancers cell lines. EBV can cause infectious mononucleosis, also called mono, and other illnesses. EBV has coevolved and become ubiquitous in all human populations through its different hosts, its ability to establish lifelong latency, intermittent reactivation after primary infection and limited clinical symptoms in the majority of infected individual[12].

The Raji cell line of lymphoblast-like cells was established from a Burkitt’s lymphoma of the left maxilla of an 11-year-old Negro male which have become invaluable tools for hematological research as they provide an unlimited amount of cellular material. Cells of the Raji line do not contain virus particles as demonstrated by electron microcopy and although the cells are resistant to vesicular stomatitis virus, this resistance is not transferred to other normally susceptible test cultures and an interferon-like inhibitor has not been found[13].

Metabolomics is broadly defined as the large-scale study of systematic identification and quantification of the small molecule metabolic products (the metabolome) of a biological system (cell, tissue, organ, biological fluid, or organism), which are the end products of cellular processes. Metabolome refers to the complete set of small-molecule metabolites (such as carbohydrate, amino acids, nucleotides, phospholipids, steroids, fatty acids, metabolic intermediates, hormones and other signaling molecules, and secondary metabolites)[14]. The metabolome forms a large network of metabolic reactions, where outputs from one enzymatic chemical reaction are inputs to other chemical reactions. Mass spectrometry and NMR spectroscopy are the techniques most often used for metabolomics profiling[15,16].

Various studies have shown that EBV have been an important cause of cancer in human and is associated with a broad-spectrum of human cancers originating from epithelial cells, lymphocytes and mesenchymal cells. Recent advances in cancer research have demonstrated that Epstein Barr virus can alter the metabolites of infected cells and may result in the development of cancer in humans[4–11]. In recent years, LC-MS and 1H NMR have been established as the gold standard technique for metabolites analysis because of the technique’s inherent analytical specificity and sensitivity[16,17]. The purpose of these efforts is for identification and quantification of metabolites that are uniquely correlated with an individual disease in order to accurately diagnose and treat the morbidity. Previous reports revealed that some metabolites (tripenoides) in raji cells may have inhibitory effects on the induction of Epstein-Barr virus early antigen (EBV-EA) by 12-O-tetradecanoylphorbol-13-acetate (TPA) which metabolites were determined on the basis of spectroscopic methods[18,19].

The purpose of precision medicine is to design disease prevention and treatment methods taking into account individual variability in environment, lifestyle, genetics, and molecular phenotype which metabolic phenotyping has the potential to generate high-volumes of complex spectral data[20]. The present study aimed to detect the altered metabolites of Raji cells in the presence of Epstein-Barr virus (EBV) by using 1H NMR Spectroscopy.

## Materials and methods

### Cell line

The Raji (B-cell lymphocyte) and EBV-producing marmoset B-cell (B95-8) cell lines were obtained from national Cell Bank of Iran (Pasteur Institute, Tehran, Iran)[21,22].

### EBV preparation

B95-8 cells were cultivated in RPMI 1640 with 15% fetal bovine serum at 37°C and in 5% CO2 humidified atmosphere. After 48 hours, the cells were centrifuged at 1300 rpm for 4 min to separate EBV-containing culture supernatant from cells[22].

### Cell culture

The Raji cells were maintained in RPMI 1640 supplemented with L-glutamine, 10% FBS and 1% antibiotics (penicillin/streptomycin). The cells (1 × 10^6^ cells/ml) were plated in T-25 flasks containing 5 ml of CGM and grown in a humidified incubator under an atmosphere of 95% air and 5% CO2 at 37°C to sub confluence (90 - 95%). The Raji cells were transferred to 6 falcon tubes and were centrifuged at 1300 rpm for 5 min[23].

### Virus transfection

Viruses were obtained from B95-8 cells. 1 ml of cells had transfected into falcon tubes containing Raji cells and incubated at 37°C for 2 hours. After incubation, 4 ml of CGM supplemented with 5% fetal bovine serum was added to each falcon tubes and the contents of the falcon tubes were transferred to a new flask. After one week, Raji cells were infected by EBV[23].

### PCR analysis

Raji cells were seeded in dishes at 500,000 cells/10 mL/ 75 cm2. One day after seeding, the medium was changed, and the cells were incubated with the test compounds for 12 h. At the end of the incubation, DNA was extracted using QIAGEN kit (QIA amp^®^ DNA Mini & Book Mini Handbook) and was refrigerated at −20°C. PCR for EBNA I gene was carried out using the specific primers which forward and reverse primers were prepared from LIGO company. The results of the PCR productshave been confirmed by an agarose gel electrophoresis.

### Cell extraction

The method of extraction using methanol-chloroform-water was done as described previously. The temperature of the extraction procedure was maintained at 4°C by working in a crushed ice bath. Cells were washed in 1X PBS and centrifuged at 6,000g for 5 min and resuspended in 500 μL of ice-cold 2:1 (v/v) methanol-chloroform solution. It was then transferred into a 1.5mL Eppendorf tube, 250 μL of ice-cold H2O 1:1 (v/v) chloroform/H20 was added and vortexed. The cells were sonicated on ice for 10min and centrifuged for 5min at 18000X*g*. The lower lipophilic and the upper hydrophilic extracts were separated and collected in different Eppendorf tubes and lyophilized and stored at −20°C [24].

### 1H NMR preparation

Lyophilized hydrophilic cell extracts (n=10) were resuspended in 200 μL of buffer (150mM potassium phosphate at pH 7.4, 1mM NaN3, and 0.01% trimethylsilyl propionate (TSP) (Sigma, CA, USA) in 100% D2O and the lipophilic cell extracts (n=10) were resuspended in 200 μLdeuterated chloroform.

### 1H NMR spectroscopy

The cell suspensions were placed in 5mm probes for analysis and one dimensional spectroscopy was performed on a 1H NMR spectrometer (Bruker AV-500) with filed gradient operating at 500.13 MHZ for observation of proton at 298K. One dimensional 1H NMR spectra were acquired with 6009.6 Hz spectral width, a 10-μs pulse 0.1 s mixing time, 3000 transients and 3.0 s relaxation delay, with standard 1D NOESY (nuclear Overhauser spectroscopy) pulse sequence to suppress the residual water peak. The 1H NMR spectrum comprising of Fourier Transformed information about metabolites in the control and treated groups(both hydrophilic and lipophilic extracts of each) were imported into MATLAB (v.7.8.0.347) software and first analyzed by ProMetab software (version 1.1) Chemical shifts between 0 and 10 ppm were normalized and spectra binned in 0.004units and the water peak (4.7) removed. The Excel files were then assessed by PLS-Toolbox version 3.0 and Partial Least Square Discriminant Analysis (PLS-DA) was applied.

### Identification of metabolites

The differentiating metabolites related to these resonances were identified by chemical shift determination using Human Metabolome Database Data Bank (HMDB) (http://www.hmdb.ca/metabolites). The metabolic cycles were determined using Metabo-Analyst software (http://www.metaboanalyst.ca/).

## Results

Figure 1 shows Superimposed 1H NMR spectra of hydrophilic and lipophilic layers between experimental and control groups.

**Fig. 1.**
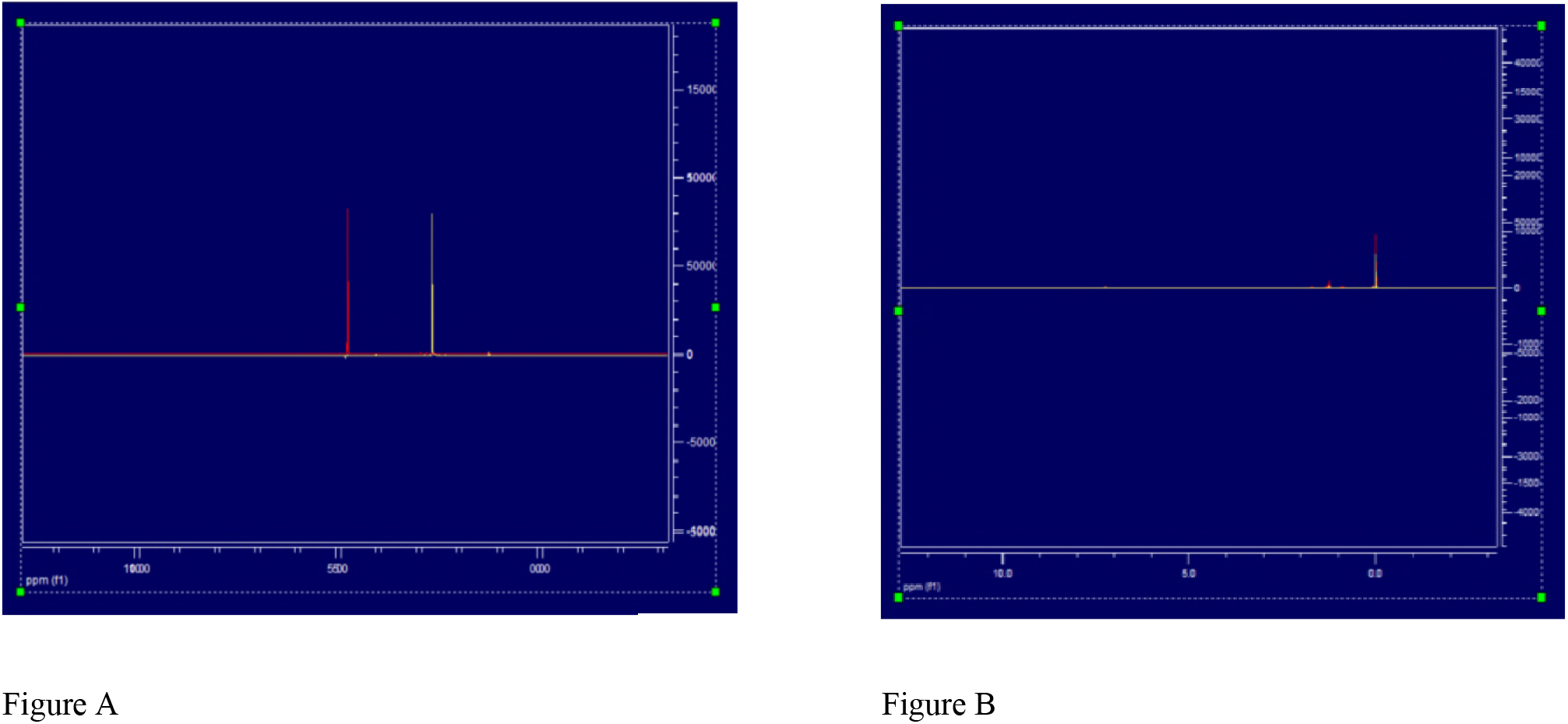
Superimposed 1HNMR spectra ofhydrophilicphase (A) and lipophilic phase (B) of EBV treated Raji cells and controls.

Figure 2 Indicates the analysis of 1H NMR spectra of hydrophilic and lipophilic layers is depicted as score plot using Partial Least Squares – Discriminant Analysis (PLS-DA) method.

**Fig. 2.**
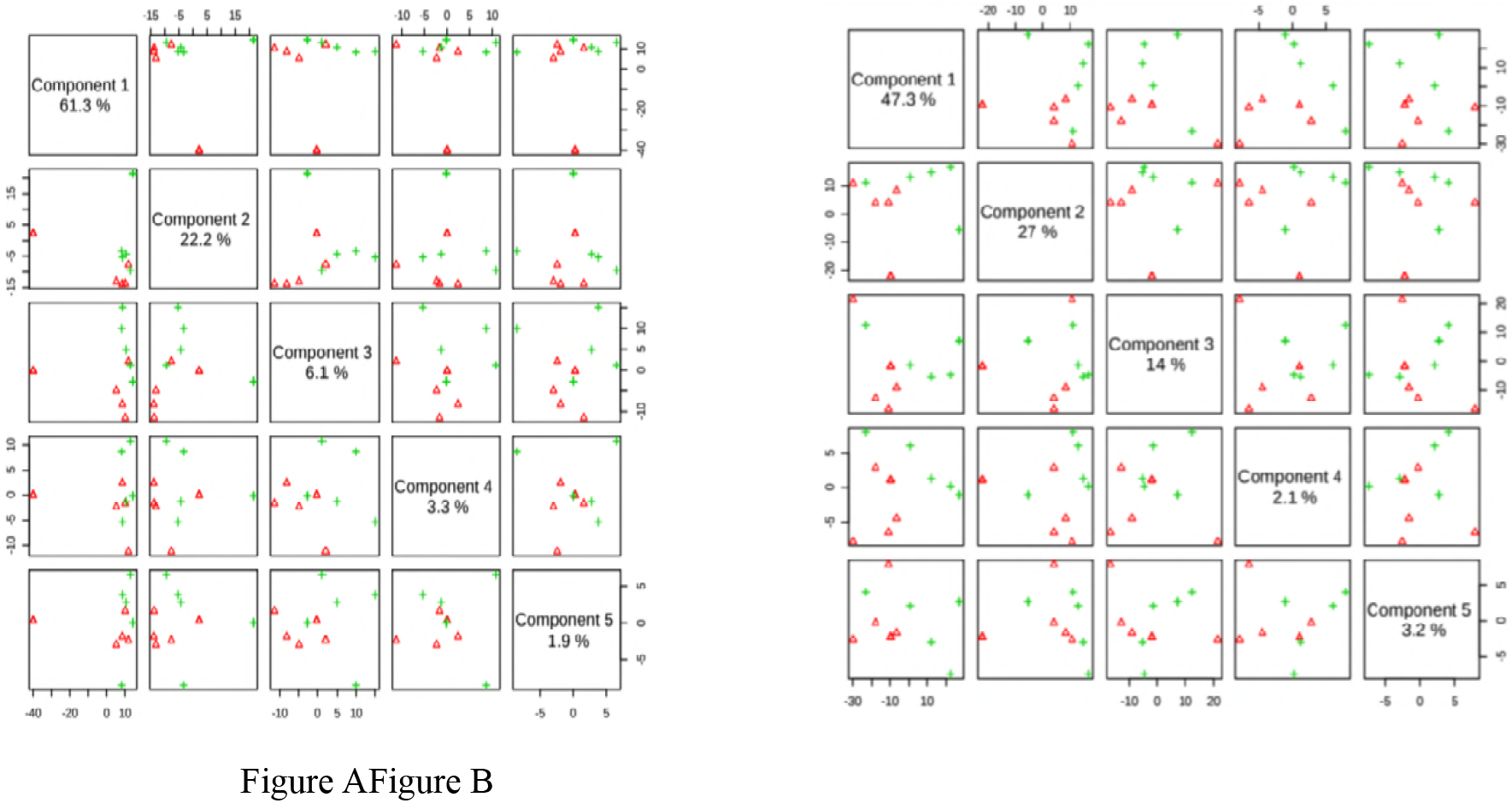
Score plot of Raji cells with EBV inexperimental and control groups inhydrophilic phase (A) and lipophilic phase (B) using PLS-DA method.

Figure 3 shows loading plot of hydrophilic and lipophilic layers between experimental and control groups using PLS-DA method.

**Fig. 3.**
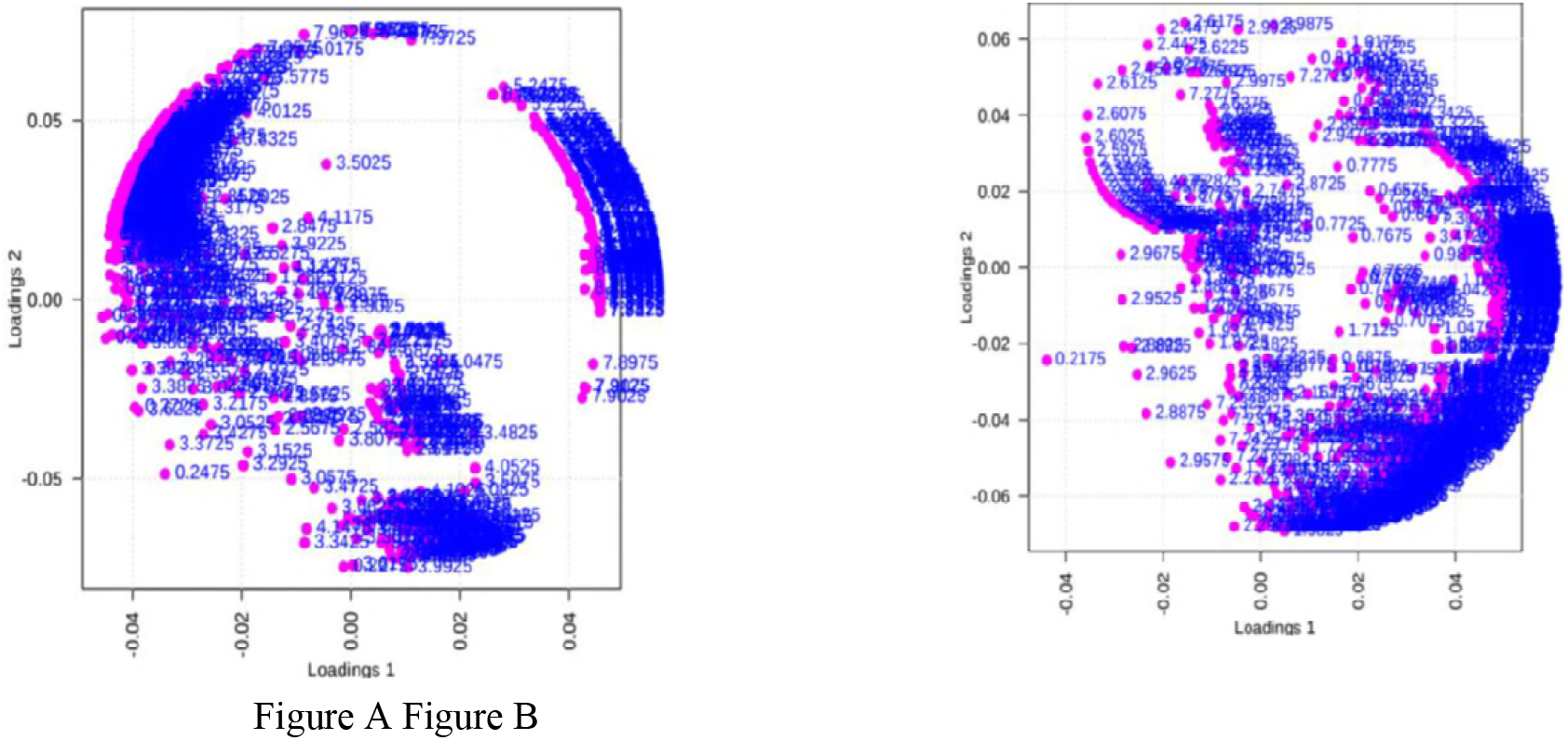
Loading plot of Raji cells with EBV inexperimental and control groups inhydrophilic phase (A) and lipophilic phase (B) using PLA-DA method.

Figure 4 shows the affected metabolites of the raji cells in hydrophilic and lipophilic phase treated by EBV using enrichment analysis.

**Fig. 4.**
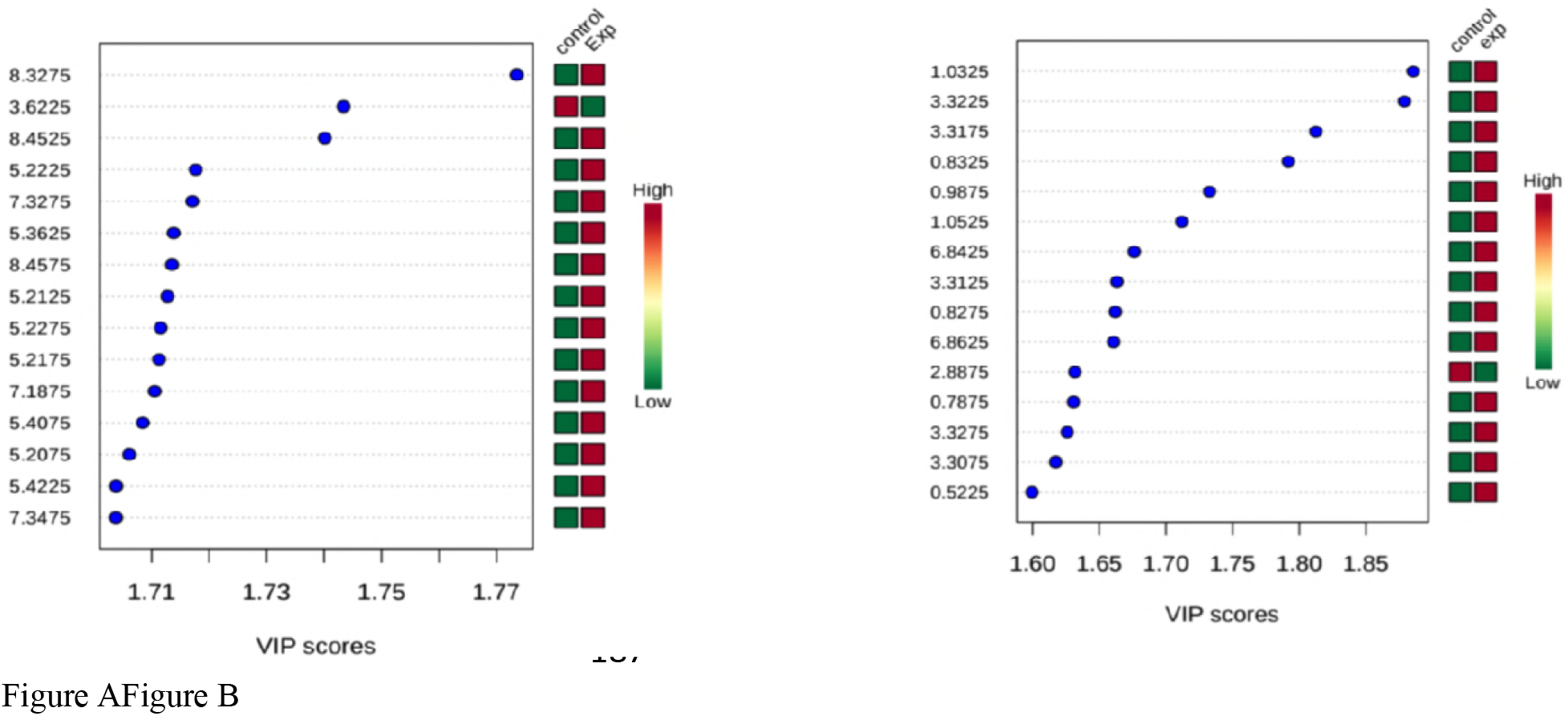
Summary plot of over-representation analysis ofhydrophilic phase (A) and lipophilic phase (B) metabolites of Raji cells treated with EBV.

Table 1 showed the results of metabolites of Raji cells in hydrophilic phase treated by EBV which obtained from human metabolites data base (HMDB).

**Table. 1.** Altered metabolites of treated group with EBV in hydrophilic phase on the base of HMDB.

| Metabolite | HMDB | Chemical Shift | levels | Abbreviation |
| --- | --- | --- | --- | --- |
| L-Phenylalanine | HMDB00159 | 7.3475 | ↑ | L-Phe |
| Glucose 1-phosphate | HMDB01586 | 5.4225 | ↑ | Glu 1-P |
| L-Fucose | HMDB00174 | 5.2075 | ↑ | L-Fu |
| D-Ribose 5-phosphate | HMDB01548 | 5.4075 | ↑ | D-Rib |
| L-Tyrosine | HMDB00158 | 7.1875 | ↑ | L-Tyr |
| L-Fucose | HMDB00174 | 5.2175 | ↑ | L-Fuc |
| Glucose 6-phosphate | HMDB01401 | 5.2275 | ↑ | Glu 6-p |
| 1-Methyladenosine | HMDB03331 | 5.2125 | ↑ | 1.Methyladenosine |
| PC(16:0/16:0) | HMDB00564 | 8.4575 | ↑ | PC |
| L-Phenylalanine | HMDB00159 | 5.3625 | ↑ | L- Phe |
| Glucose 6-phosphate | HMDB01401 | 7.3275 | ↑ | Glu 6-p |
| D-Ribose 5-phosphate | HMDB01548 | 5.2225 | ↑ | D-Rib 5-p |
| 1-Methyladenosine | HMDB03331 | 8.4525 | ↑ | 1.Methyladenosine |
| Fructose 6-phosphate | HMDB00124 | 3.6225 | ↓ | Fru 6-p |
| Nicotinamideribotide | HMDB00229 | 8.3275 | ↑ | Nicotinamideribotide |

Demonstration of the alteration of metabolites of Raji cells in lipophilic phase treated by EBV which obtained from human metabolites data base (HMDB) showed in Table 2.

**Table. 2.** Altered metabolites of treated group with EBV in lipophilic phase on the base of HMDB.

| Metabolite | HMDB | Chemical Shift | levels | Abbreviation |
| --- | --- | --- | --- | --- |
| Cholesterol | HMDB00067 | 0.5225 | ↑ | Chol |
| L-proline | HMDB00162 | 3.3075 | ↑ | L-Pro |
| Beta-Leucine | HMDB03640 | 3.3275 | ↑ | β- Ala |
| Cholesterol | HMDB00067 | 0.7875 | ↑ | Chol |
| L-Arginine | HMDB0000168 | 2.8875 | ↓ | L-Arginine |
| 5-Hydroxy-L-tryptophan | HMDB00472 | 6.8625 | ↑ | 5-Hydroxy-L-Try |
| Tryptophan | HMDB00929 | 6.8625 | ↑ | Try |
| Cholesterol sulfate | HMDB0000653 | 0.8275 | ↑ | Chol |
| L-Proline | HMDB00162 | 3.3125 | ↑ | L-Pro |
| S-Adenosylhomocysteine | HMDB00939 | 6.8425 | ↑ | SAH-Cys |
| 2-Ketobutyric acid | HMDB00005 | 1.0525 | ↑ | 2- Ketobutyric acid |
| L-Valine | HMDB00883 | 0.9875 | ↑ | L-Val |
| Cholesterol | HMDB00067 | 0.8325 | ↑ | Chol |
| L-Proline | HMDB00162 | 3.3175 | ↑ | L-Pro |
| L-Proline | HMDB00162 | 3.3225 | ↑ | L-Pro |
| L-Valine | HMDB00883 | 1.0325 | ↑ | L-Val |

Figure 5 shows the affected metabolic pathways of the raji cells in hydrophilic phase treated by EBV using the pathway analyzing tool (MetaboAnalyst 3.0).

**Fig. 5.**
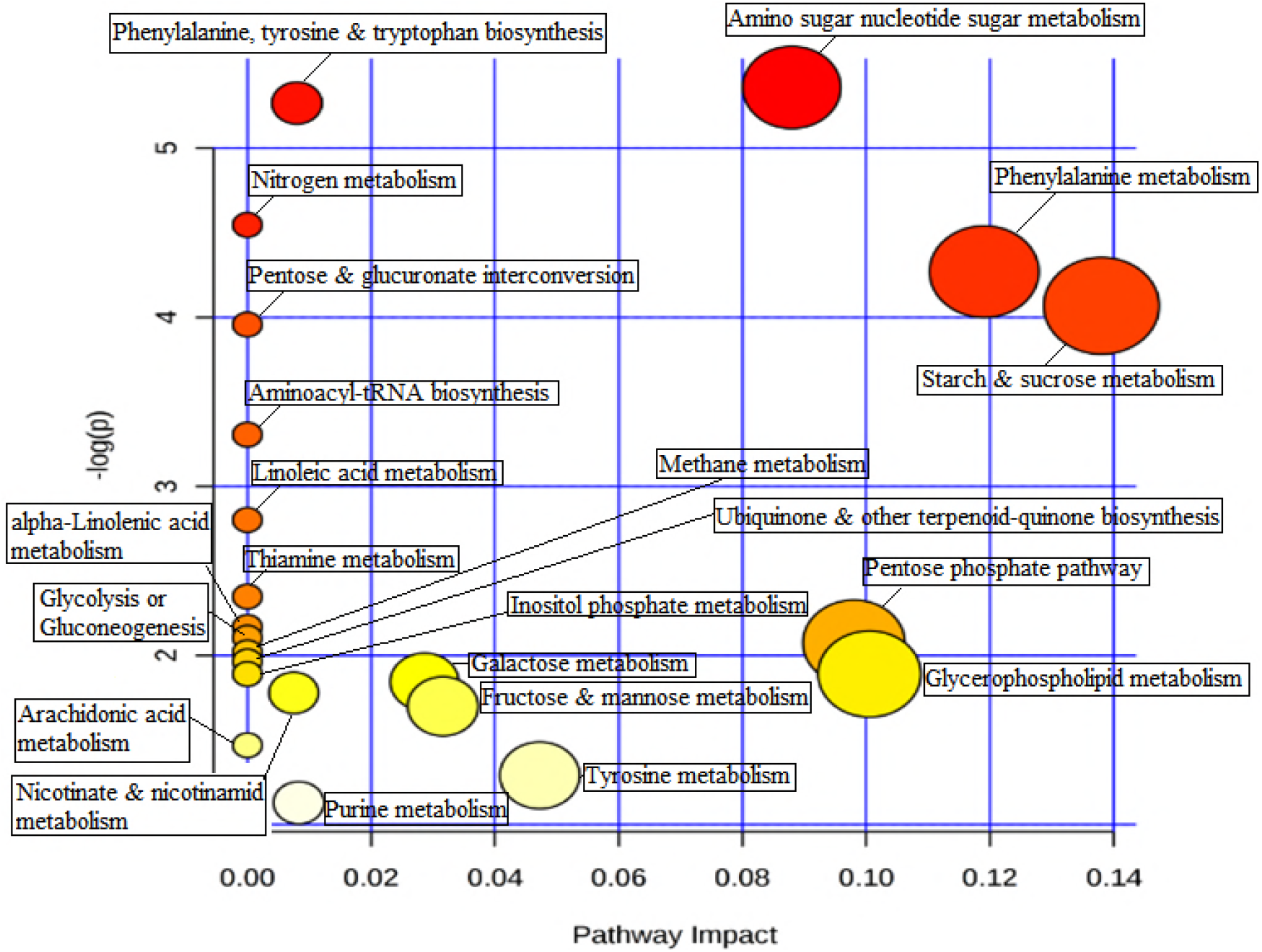
Important metabolicpathways in the hydrophilic of Raji cells treated with EBV using MetaboAnalyst 3.0.

Table 3 indicates the alteration of metabolic pathways of Raji cells in hydrophilic phase treated by EBV using the pathway analyzing tool (MetaboAnalyst 3.0).

**Table. 3.**
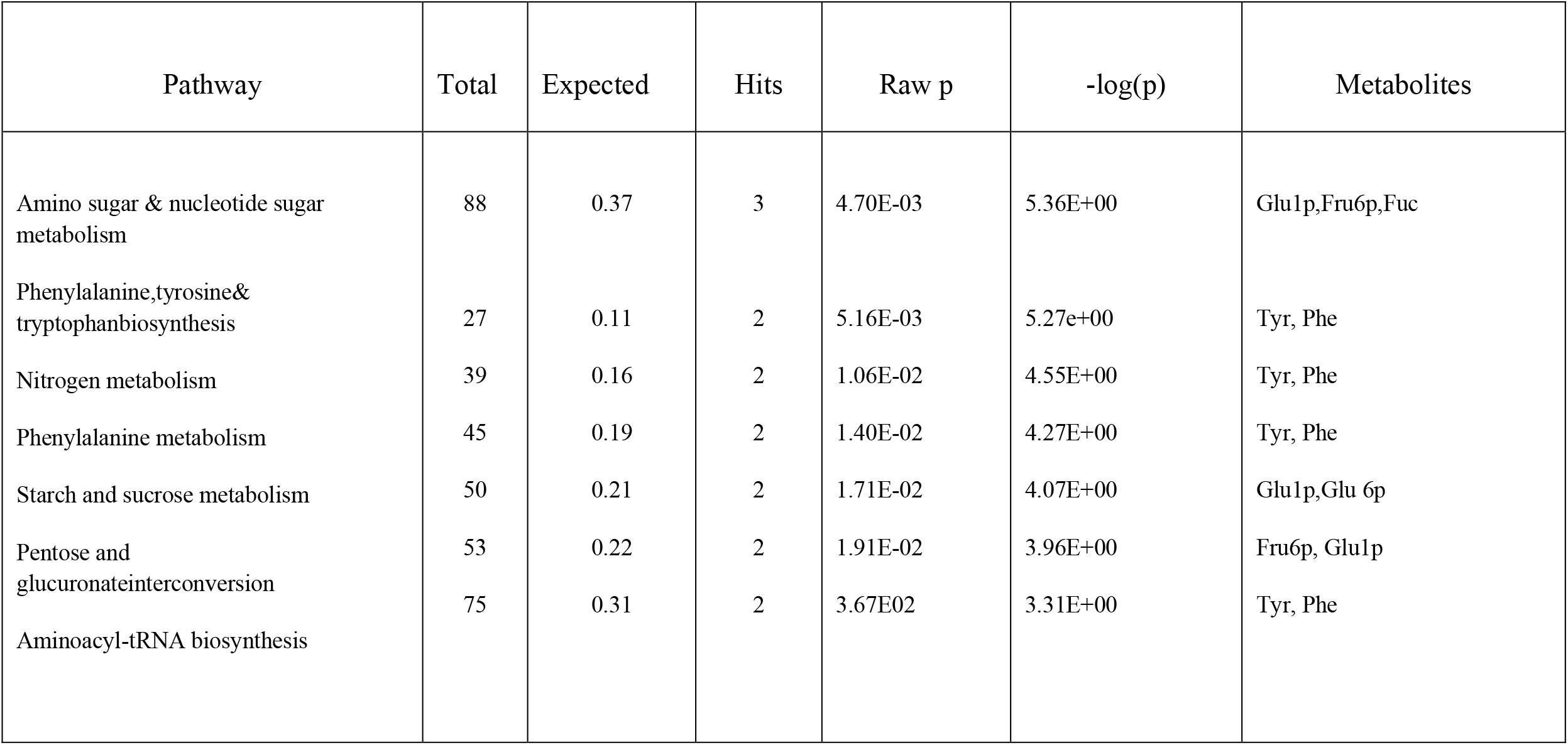
Altered metabolic pathways of treated group with EBV in hydrophilic phase using MetaboAnalyst 3.0.

Figure 6 indicates the affected metabolic pathways of the raji cells in lipophilic phase treated by EBV using the pathway analyzing tool (MetaboAnalyst 3.0).

**Fig. 6.**
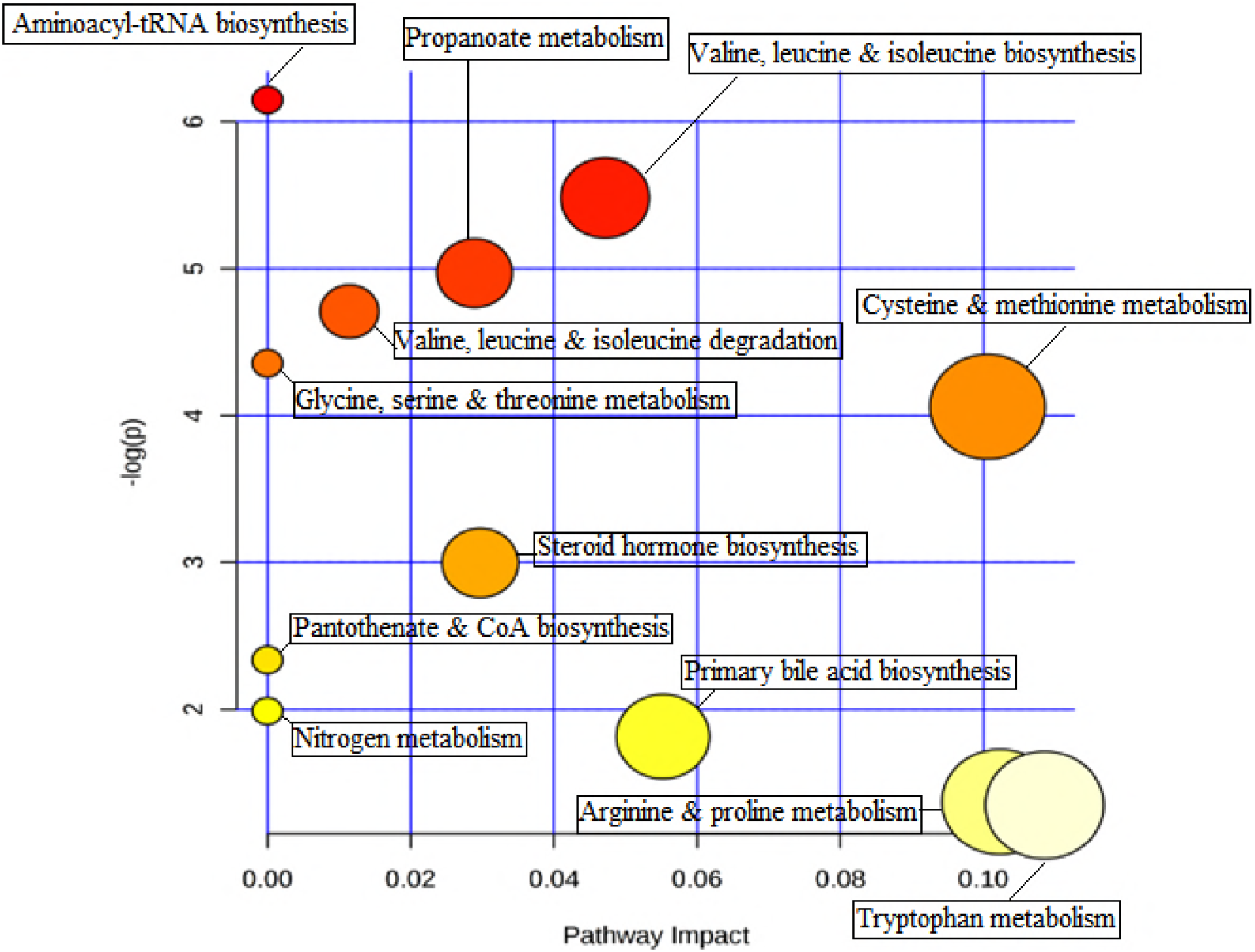
Important metabolicpathways in the lipophilic phaseofRaji cells treated with EBV using MetaboAnalyst 3.0.

Table 4 demonstrates the alteration of metabolic pathways of Raji cells in lipophilic phase treated by EBV using the pathway analyzing tool (MetaboAnalyst 3.0).

**Table 4.** ....................

| Pathway | Total | Expected | Hits | Raw p | -log(p) | Metabolites |
| --- | --- | --- | --- | --- | --- | --- |
| Aminoacyl-tRNA biosynthesis | 75 | 0.28 | 3 | 2.31E-03 | 6.15E+00 | Pro, Tyr, tRNA(Val) |
| Valine, leucine & isoleucine biosynthesis | 27 | 0.1 | 2 | 4.16E-03 | 5.48E+00 | Val, Ketobutyric acid |
| Propanoate metabolism | 35 | 0.13 | 2 | 6.94E-03 | 4.97E+00 | Val, Ketobutyric acid |
|  | 40 | 0.15 | 2 | 9.01E-03 | 4.71E+00 | Val, Leu |
| Valine, leucine and isoleucine degradation | 48 | 0.18 | 2 | 1.28E-02 | 4.36E+00 |  |
| Glycine, serine & threonine metabolism | 99 | 0.37 | 2 | 4.99E-02 | 3.00E+00 | Trp, Ketobutyric acid |
| Steroid hormone biosynthesis |  |  |  |  |  | Chol, L-Arg |

## Discussion

A wide range of metabolites and chemical structure were measured upon extraction and samples preparation, which berecognizedsignificantly, altered metabolic pathways according to metaboAnalyst website.

According to the obtained results for hydrophilic phase, glucose 1-phosphate, glucose 6- phosphate and fructose 6-phosphate have been the most frequency among metabolic pathways including amino sugar and nucleotide sugar reaction, starch and sucrose metabolism and pentose-glucuronateinterconversion. In lipophilic phase, proline, valine, tyrosine, triptophane, and ketobutyric acid also have the most alteration among metabolic pathway including amino acyl tRNA, valine, leucine and isoleucine biosynthesis, glycine, serine, threonine and propaonate metabolism and valine, leucine and isoleucine degradation aromatic amino acids ( tyrosine and phenylalanine ) also may be changed among tyrosine, phenylalanine, tryptophane and nitrogen metabolic pathways. Although,the alteration in metabolic pathways have been resulted in difference between Raji cells and EBV-infected cells.

The results indicated that amino sugar and nucleotide sugar metabolism are the most important metabolic pathways in hydrophilic phase. The metabolites conversions, especially glucose 1-phosphate,fructose 6-phosphate and fucose were observed in these pathways, which they are the most valuable metabolites in glycolysis pathway. The result from this study also support that changes in metabolite concentration, particularly rising glucose and L-fucose levels were found in EBV-infected raji cells in comparison with control cells. It also hasbeenshown that fructose 6- phosphate and glucose 1-phosphateare involved in pentose-glucuronateinterconversion pathway. In line with this investigation, some virological studies showed that EBV lead to change in metabolic assay from early infection to long-term outgrowth that may be stimulate glucose import and surface glucose transporter-1 (Glut-1) levels, result in induction of glycolysis, oxidative phosphorylation and suppression of basal autophagy[25]. There are some reports showing that serum glycoprotein L-fucose levels have two-fold rise in head and neck neoplasma compared to control group[26]. Advanced in cancer have been shown that fucose may be is useful in breast cancer treatments and α-L-fucose has pivotal role in construction of malignant and metastaric phenotype of various human breast cancer cell lines. Additionally, some breast cancer cell lines biomarkers are fucose-rich[27].

Metabolomics approach demonstrate alteration in glycolysis metabolites is associated with EBV in nasopharyngeal carcinoma (NPC) which overexpression of EBV-encoded latent protein 1 (LMP1) may lead to glycolysis induction. Some glycolysis genes (i.e. hexokinase 2) have central roles in LMP1-mediated glucose metabolism reprogramming in NPC cells. Additionally, positive correlation was existed between HK2 and LMP1 in NPC biopsies, and the HK2 induction was associated with poor survival of NPC patient after radiation therapies[28]. Therefore, there is a potential correlation between glycolysis pathway and EBV infected cells. Furthermore, the pentos phosphate pathway is required for ribonocleotide synthesis and NADPH production which branches from glycolysis [29].At first this research suggests phenylalanine and tyrosine are involved in nitrogen metabolism pathway. According to virologycal studies, two phenylalanine (F600, F605) are located in R transactivator (Rta) c-terminal which play pivotal role in DNA binding to target cells. These two phenylalanine are essential for Rta expression which Rta activates the EBV lytic cycle. If two other aromatic amino acids (Tryptophan and Tyrosine) are substituted with two phenylalanine, maintenance of mRNA activity of the BMLF1 gene have been seen. However, substitution of Tryptophan and Tyrosine with non-aromatic amino acid including Alanine and Valine, lead to capacity elimination of Rta activity. Valine and glycine substitution instead of phenylalanine in Rta protein act as inhibitor and may prevent its DNA-binding function. The EBV BZLF1 protein (ZEBRA, Zta) aromatic amino acids including (phenylalanine, Tyrosine and Tryptophan) are crucial components of activation domain,thereforeRta play important role in EBV lytic cycle by the subsitution of cellular signaling pathway and synergy with EBV ZEBRA protein [30].

Moreover, the results in lipophilic phase revealed that proline, valine, tyrosine, tryptophane amino acids and ketobutyric acid are altered in metabolic pathway of tRNA, valine, leucine, isoleucine, serine, glycine, threonine and propanoate biosynthesis, and degradation of valine, leucine and isoleucine. Amino acyletRNA biosynthesis could be among the first pathways in lipophilic phase which several amino acids such as proline, tyrosine and valine can be altered in thesepathways. N-terminal domain of LMP2A contains eight phosphorylated tyrosine residue.Two of which constitute an immunoreceptor tyrosine activation motif (ITAM)[31]. ITAM consist of a pared tyrosine and leucine residues and play a pivotal role in signal transduction of B-cell receptors (BCR) and T-cell receptors, lymphocyte proliferation and activity of kinase families. Although, LMP2A ITAM motif participate in BCR signal transduction as an inhibitor[32]. Several lines of studies suggest that LMP2A prevents BCR signal transduction trough engaging Nedd-4 ubiquitin protein ligases to promote the degradation of Lyne and LMP2A by an ubiquitin-dependent mechanism[33].

The results showed in lipophilic phase also indicating that valine and acid butyric are involved and altered in metabolic pathway of valine, leucine and isoleucine biosynthesis. Several investigation consider that the C-myc transcription factor is acting as a proto-oncogene which activation of MYC lead to promotion of cell cycle transition and is recognized as a leucine zipper protein which is activation by mitogenic factors under normal circumstances[24]. On this base, previous studies on EBV genome have shown that multi-nucleonal proteins have a highly charged N-terminus which may provide nuclear signals and contains heptad repeats of leucine, isoleucine or valine that can act as dimerization domain; the third exon includes leucine and isoleucine heptad repeats which make possible intraction of coiled-coild and facilities in hemodimerization of BZLF1[35].

In present study tryptophane level has dramatically increased in glycine, serine and threonine metabolic pathways. Valine and ketobutyric acid also have seen to be engaged in propaonate metabolism’srecent data, indicating that Tryptophane is essential for virus penetration in cells and crosses lipid bilayers without pore formation[36].

In steroid hormone metabolism, the most alteration was observed in amount of cholesterol metabolites. Some investigation has revealed that cholesterol enriched-domains in plasma membranes may are required for infection of human B-cells with EBV and is necessary for membrane fusion, receptor localization in membrane micro-domain and early viral signaling events. Lipid rafts are also involved in MHC class II protein function[37].

Additionally, the finding results demonstrate that L-Argenine level has significantly reduced in cells exposed to EBV compared to control cells. There is various reports indicating L-Argenine supplementation may be lead to inhibition of spontaneous EBV reactivation in another Burkitts lymphoma cell line EB1 and B lymphoblastic cell line OB. L-Argenine also can induces inducible NO synthase and generates NO, which inhibits EBV reactivation in EBV-possitive cells[38].

The conclusionof this research showed that infection of Rajicells with EBV leads to proliferation and subsequent immortalization of cell lines through cellular replication machinery recruitment andchanges the metabolic profile and promotes vital metabolites such as the carbohydrates (i.e. glucose-1-phosphate and fructose 1-phosphate) engage in pentose phosphate and glycolactic, biosynthesis of nucleotide and amino acid pathways. Theresults also indicate that essential amino acids are required for protecting viral structure and the function of viral genes. Furthermore, rates of proteins synthesis and function of glycolysis pathway give rise to increase in EBV-infected Raji cells compare with control cells. Therefore, EBV infection of Raji cells leads to the sustained elevation of cell growth and cell immortalization.

## Acknowledgement

I would like to express my thanks to Pasteur Institute of Iran’s Research council members especially Dr. M.R Aghasadeghi as the head of Hepatitis & AIDS & also staf of Pasteur Institute of Iran. Finally, I would like to express my appreciation to Dr. Hajhosseini as the head of Payam Noor University of Tehran for his supports.

